## Supplementary file for "Drivers of species knowledge across the Tree of Life"

**Supplementary File 1a. Estimated regression parameters for the full models modeling the influence of species-level traits and cultural factors on scientific (Web of Science) and societal interest (Wikipedia) for different taxa, as well as the relative scientific and societal interest (Residuals).** CI: 95% Confidence interval.

| Model | Type | Parameter | Beta | SE | CI_low | CI_high | z | p |
|---|---|---|---|---|---|---|---|---|
| Web of Science | - | Intercept | 0.889 | 0.492 | -0.075 | 1.853 | 1.808 | 0.071 |
| Web of Science | Species trait | Organism size | 0.445 | 0.111 | 0.228 | 0.663 | 4.012 | <0.001 |
| Web of Science | Species trait | Colorful [yes] | 0.217 | 0.145 | -0.068 | 0.502 | 1.493 | 0.136 |
| Web of Science | Species trait | Range size | 0.598 | 0.057 | 0.486 | 0.71 | 10.481 | <0.001 |
| Web of Science | Species trait | Domain [freshwater] | 0.251 | 0.315 | -0.366 | 0.868 | 0.797 | 0.426 |
| Web of Science | Species trait | Domain [marine] | -0.27 | 0.279 | -0.817 | 0.276 | -0.969 | 0.332 |
| Web of Science | Species trait | Domain [terrestrial] | -0.546 | 0.216 | -0.97 | -0.122 | -2.525 | 0.012 |
| Web of Science | Species trait | Taxonomic uniqueness (Family) | -0.408 | 0.077 | -0.559 | -0.257 | -5.306 | <0.001 |
| Web of Science | Cultural trait | IUCN [threatened] | 1.229 | 0.192 | 0.853 | 1.604 | 6.41 | <0.001 |
| Web of Science | Cultural trait | IUCN [non-threatened] | 0.974 | 0.154 | 0.672 | 1.277 | 6.315 | <0.001 |
| Web of Science | Cultural trait | Common name [yes] | 0.968 | 0.165 | 0.645 | 1.291 | 5.874 | <0.001 |
| Web of Science | Cultural trait | Human use [yes] | 1.026 | 0.121 | 0.788 | 1.264 | 8.456 | <0.001 |
| Web of Science | Cultural trait | Harmful to humans [yes] | 1.732 | 0.243 | 1.257 | 2.208 | 7.139 | <0.001 |
| Web of Science | Cultural trait | Phylogenetic distance to humans | -0.73 | 0.382 | -1.479 | 0.02 | -1.907 | 0.056 |
| Wikipedia | - | Intercept | 5.973 | 0.354 | 5.28 | 6.666 | 16.891 | <0.001 |
| Wikipedia | Species trait | Organism size | 0.844 | 0.096 | 0.657 | 1.032 | 8.841 | <0.001 |
| Wikipedia | Species trait | Colorful [yes] | 0.744 | 0.125 | 0.498 | 0.99 | 5.932 | <0.001 |
| Wikipedia | Species trait | Range size | 0.214 | 0.037 | 0.14 | 0.287 | 5.701 | <0.001 |
| Wikipedia | Species trait | Domain [freshwater] | 0.827 | 0.291 | 0.256 | 1.398 | 2.838 | 0.005 |
| Wikipedia | Species trait | Domain [marine] | 0.3 | 0.287 | -0.261 | 0.862 | 1.048 | 0.295 |
| Wikipedia | Species trait | Domain [terrestrial] | 0.248 | 0.214 | -0.172 | 0.668 | 1.157 | 0.247 |
| Wikipedia | Species trait | Taxonomic uniqueness (Family) | -0.416 | 0.067 | -0.548 | -0.284 | -6.186 | <0.001 |
| Wikipedia | Cultural trait | IUCN [threatened] | 0.841 | 0.162 | 0.522 | 1.159 | 5.176 | <0.001 |
| Wikipedia | Cultural trait | IUCN [non-threatened] | 0.303 | 0.126 | 0.055 | 0.55 | 2.397 | 0.017 |

| | | | | | | | | |
|---|---|---|---|---|---|---|---|---|
| Wikipedia | Cultural trait | Common name [yes] | 1.491 | 0.15 | 1.196 | 1.785 | 9.919 | <0.001 |
| Wikipedia | Cultural trait | Human use [yes] | 0.985 | 0.109 | 0.771 | 1.199 | 9.028 | <0.001 |
| Wikipedia | Cultural trait | Harmful to humans [yes] | 1.827 | 0.242 | 1.354 | 2.301 | 7.56 | <0.001 |
| Wikipedia | Cultural trait | Phylogenetic distance to humans | -0.957 | 0.287 | -1.519 | -0.395 | -3.337 | 0.001 |
| Residuals | - | Intercept | -2.291 | 0.41 | -3.095 | -1.487 | -5.584 | <0.001 |
| Residuals | Species trait | Organism size | 0.276 | 0.103 | 0.074 | 0.478 | 2.684 | 0.007 |
| Residuals | Species trait | Colorful [yes] | 0.426 | 0.133 | 0.165 | 0.688 | 3.197 | 0.001 |
| Residuals | Species trait | Range size | -0.162 | 0.046 | -0.253 | -0.072 | -3.527 | <0.001 |
| Residuals | Species trait | Domain [freshwater] | 0.042 | 0.332 | -0.607 | 0.692 | 0.128 | 0.898 |
| Residuals | Species trait | Domain [marine] | 0.637 | 0.309 | 0.031 | 1.243 | 2.06 | 0.039 |
| Residuals | Species trait | Domain [terrestrial] | 0.464 | 0.223 | 0.028 | 0.9 | 2.084 | 0.037 |
| Residuals | Species trait | Taxonomic uniqueness (Family) | -0.187 | 0.071 | -0.327 | -0.047 | -2.614 | 0.009 |
| Residuals | Cultural trait | IUCN [threatened] | -0.179 | 0.195 | -0.561 | 0.204 | -0.915 | 0.36 |
| Residuals | Cultural trait | IUCN [non-threatened] | -0.334 | 0.153 | -0.634 | -0.033 | -2.179 | 0.029 |
| Residuals | Cultural trait | Common name [yes] | 0.71 | 0.155 | 0.406 | 1.013 | 4.584 | <0.001 |
| Residuals | Cultural trait | Human use [yes] | -0.312 | 0.129 | -0.565 | -0.059 | -2.418 | 0.016 |
| Residuals | Cultural trait | Harmful to humans [yes] | -0.32 | 0.269 | -0.847 | 0.207 | -1.189 | 0.234 |
| Residuals | Cultural trait | Phylogenetic distance to humans | -0.475 | 0.304 | -1.072 | 0.121 | -1.561 | 0.118 |

**Supplementary File 1b. Estimated regression parameters for the subset models modeling the influence of species-level traits and cultural factors on scientific interest (N° opf papers in the Web of Science) for Chordata, Arthropoda and Tracheophyta.** CI: 95% Confidence interval; NA: Not Available.

| **Model** | **Type** | **Parameter** | **Beta** | **SE** | **CI_low** | **CI_high** | **z** | **p** |
|---|---|---|---|---|---|---|---|---|
| Chordata | Intercept | Intercept | -0.144 | 0.649 | -1.415 | 1.128 | -0.221 | 0.825 |
| Chordata | Species trait | Organism size | 1.423 | 0.207 | 1.016 | 1.829 | 6.858 | <0.001 |
| Chordata | Species trait | Colorful [yes] | 0.238 | 0.205 | -0.163 | 0.639 | 1.163 | 0.245 |
| Chordata | Species trait | Range size | 0.818 | 0.085 | 0.65 | 0.985 | 9.567 | <0.001 |
| Chordata | Species trait | Domain [freshwater] | 0.315 | 0.451 | -0.57 | 1.199 | 0.697 | 0.486 |
| Chordata | Species trait | Domain [marine] | -0.797 | 0.387 | -1.556 | -0.038 | -2.057 | 0.04 |
| Chordata | Species trait | Domain [terrestrial] | -0.398 | 0.223 | -0.834 | 0.039 | -1.786 | 0.074 |
| Chordata | Cultural trait | IUCN [threatened] | 1.176 | 0.202 | 0.781 | 1.571 | 5.834 | <0.001 |
| Chordata | Cultural trait | IUCN [non-threatened] | 0.891 | 0.166 | 0.565 | 1.216 | 5.363 | <0.001 |
| Chordata | Species trait | Taxonomic uniqueness (Genus) | -0.17 | 0.086 | -0.338 | -0.001 | -1.969 | 0.049 |
| Chordata | Cultural trait | Common name [yes] | 0.773 | 0.48 | -0.168 | 1.713 | 1.61 | 0.107 |
| Chordata | Cultural trait | Human use [yes] | 0.501 | 0.132 | 0.242 | 0.76 | 3.796 | <0.001 |
| Chordata | Cultural trait | Harmful to humans [yes] | 1.247 | 0.347 | 0.568 | 1.927 | 3.598 | <0.001 |
| Arthropoda | Intercept | Intercept | -0.308 | 0.97 | -2.209 | 1.592 | -0.318 | 0.75 |
| Arthropoda | Species trait | Organism size | -0.403 | 0.247 | -0.888 | 0.082 | -1.628 | 0.103 |
| Arthropoda | Species trait | Colorful [yes] | 0.439 | 0.308 | -0.165 | 1.043 | 1.423 | 0.155 |
| Arthropoda | Species trait | Range size | 0.429 | 0.111 | 0.212 | 0.646 | 3.878 | <0.001 |
| Arthropoda | Species trait | Domain [freshwater] | -0.378 | 0.978 | -2.294 | 1.539 | -0.386 | 0.699 |
| Arthropoda | Species trait | Domain [marine] | -0.48 | 0.933 | -2.308 | 1.349 | -0.514 | 0.607 |
| Arthropoda | Species trait | Domain [terrestrial] | -0.588 | 0.889 | -2.331 | 1.154 | -0.662 | 0.508 |
| Arthropoda | Cultural trait | IUCN [threatened] | 1.828 | 1.188 | -0.501 | 4.157 | 1.539 | 0.124 |
| Arthropoda | Cultural trait | IUCN [non-threatened] | -1.02 | 0.637 | -2.27 | 0.229 | -1.6 | 0.109 |
| Arthropoda | Species trait | Taxonomic uniqueness (Genus) | -0.132 | 0.109 | -0.345 | 0.082 | -1.208 | 0.227 |
| Arthropoda | Cultural trait | Common name [yes] | 0.904 | 0.27 | 0.375 | 1.433 | 3.349 | 0.001 |
| Arthropoda | Cultural trait | Human use [yes] | 3.334 | 0.646 | 2.068 | 4.6 | 5.163 | <0.001 |
| Arthropoda | Cultural trait | Harmful to humans [yes] | 2.065 | 0.442 | 1.199 | 2.932 | 4.671 | <0.001 |
| Tracheophyta | Intercept | Intercept | 0.387 | 2.114 | -3.756 | 4.53 | 0.183 | 0.855 |
| Tracheophyta | Species trait | Organism size | 0.422 | 0.285 | -0.136 | 0.981 | 1.481 | 0.139 |
| Tracheophyta | Species trait | Colorful [yes] | 0.401 | 0.374 | -0.331 | 1.133 | 1.074 | 0.283 |
| Tracheophyta | Species trait | Range size | 0.968 | 0.279 | 0.421 | 1.514 | 3.471 | 0.001 |
| Tracheophyta | Species trait | Domain [freshwater] | NA | NA | NA | NA | NA | NA |
| Tracheophyta | Species trait | Domain [marine] | NA | NA | NA | NA | NA | NA |
| Tracheophyta | Species trait | Domain [terrestrial] | -1.786 | 1.793 | -5.301 | 1.728 | -0.996 | 0.319 |

| | | | | | | | | |
|---|---|---|---|---|---|---|---|---|
| Tracheophyta | Cultural trait | IUCN [threatened] | 2.05 | 0.902 | 0.283 | 3.817 | 2.274 | 0.023 |
| Tracheophyta | Cultural trait | IUCN [non-threatened] | 1.694 | 0.558 | 0.601 | 2.788 | 3.037 | 0.002 |
| Tracheophyta | Species trait | Taxonomic uniqueness (Genus) | -0.172 | 0.15 | -0.466 | 0.121 | -1.15 | 0.25 |
| Tracheophyta | Cultural trait | Common name [yes] | 1.078 | 0.48 | 0.137 | 2.019 | 2.245 | 0.025 |
| Tracheophyta | Cultural trait | Human use [yes] | 1.047 | 0.426 | 0.211 | 1.882 | 2.455 | 0.014 |
| Tracheophyta | Cultural trait | Harmful to humans [yes] | 0.849 | 0.829 | -0.775 | 2.473 | 1.025 | 0.305 |

**Supplementary File 1c. Estimated regression parameters for the subset models modeling the influence of species-level traits and cultural factors on popular interest (views in Wikipedia) for Chordata, Arthropoda and Tracheophyta.** CI: 95% Confidence interval; NA: Not Available.

| Model | Type | Parameter | Beta | SE | CI_low | CI_high | z | p |
|---|---|---|---|---|---|---|---|---|
| Chordata | Intercept | Intercept | 7.523 | 0.701 | 6.149 | 8.898 | 10.73 | <0.001 |
| Chordata | Species trait | Organism size | 1.555 | 0.171 | 1.22 | 1.891 | 9.098 | <0.001 |
| Chordata | Species trait | Colorful [yes] | 0.446 | 0.18 | 0.093 | 0.798 | 2.479 | 0.013 |
| Chordata | Species trait | Range size | 0.446 | 0.055 | 0.339 | 0.554 | 8.118 | <0.001 |
| Chordata | Species trait | Domain [freshwater] | 2.107 | 0.393 | 1.337 | 2.877 | 5.362 | <0.001 |
| Chordata | Species trait | Domain [marine] | 0.398 | 0.412 | -0.41 | 1.207 | 0.966 | 0.334 |
| Chordata | Species trait | Domain [terrestrial] | 0.091 | 0.203 | -0.306 | 0.488 | 0.448 | 0.654 |
| Chordata | Cultural trait | IUCN [threatened] | 0.539 | 0.16 | 0.225 | 0.853 | 3.364 | 0.001 |
| Chordata | Cultural trait | IUCN [non-threatened] | -0.011 | 0.131 | -0.268 | 0.246 | -0.086 | 0.931 |
| Chordata | Species trait | Taxonomic uniqueness (Genus) | -0.261 | 0.07 | -0.398 | -0.124 | -3.742 | <0.001 |
| Chordata | Cultural trait | Common name [yes] | 0.623 | 0.34 | -0.043 | 1.29 | 1.832 | 0.067 |
| Chordata | Cultural trait | Human use [yes] | 0.645 | 0.111 | 0.428 | 0.862 | 5.826 | <0.001 |
| Chordata | Cultural trait | Harmful to humans [yes] | 1.205 | 0.328 | 0.563 | 1.847 | 3.678 | <0.001 |
| Arthropoda | Intercept | Intercept | 6.06 | 0.808 | 4.476 | 7.645 | 7.497 | <0.001 |
| Arthropoda | Species trait | Organism size | 0.857 | 0.214 | 0.437 | 1.277 | 4.002 | <0.001 |
| Arthropoda | Species trait | Colorful [yes] | 0.754 | 0.232 | 0.299 | 1.209 | 3.249 | 0.001 |
| Arthropoda | Species trait | Range size | 0.099 | 0.071 | -0.04 | 0.238 | 1.396 | 0.163 |
| Arthropoda | Species trait | Domain [freshwater] | -0.254 | 0.76 | -1.744 | 1.236 | -0.334 | 0.738 |
| Arthropoda | Species trait | Domain [marine] | -0.23 | 0.761 | -1.721 | 1.261 | -0.302 | 0.762 |
| Arthropoda | Species trait | Domain [terrestrial] | -0.009 | 0.679 | -1.34 | 1.322 | -0.013 | 0.99 |
| Arthropoda | Cultural trait | IUCN [threatened] | 1.81 | 0.995 | -0.139 | 3.76 | 1.82 | 0.069 |
| Arthropoda | Cultural trait | IUCN [non-threatened] | -0.32 | 0.571 | -1.44 | 0.8 | -0.56 | 0.575 |
| Arthropoda | Species trait | Taxonomic uniqueness (Genus) | -0.149 | 0.073 | -0.293 | -0.006 | -2.04 | 0.041 |
| Arthropoda | Cultural trait | Common name [yes] | 1.2 | 0.205 | 0.797 | 1.602 | 5.847 | <0.001 |
| Arthropoda | Cultural trait | Human use [yes] | 3.021 | 0.631 | 1.784 | 4.259 | 4.785 | <0.001 |

| | | | | | | | | |
|---|---|---|---|---|---|---|---|---|
| Arthropoda | Cultural trait | Harmful to humans [yes] | 2.357 | 0.384 | 1.605 | 3.109 | 6.141 | <0.001 |
| Tracheophyta | Intercept | Intercept | 4.479 | 1.328 | 1.876 | 7.082 | 3.373 | 0.001 |
| Tracheophyta | Species trait | Organism size | 0.268 | 0.187 | -0.098 | 0.634 | 1.436 | 0.151 |
| Tracheophyta | Species trait | Colorful [yes] | 0.495 | 0.244 | 0.017 | 0.974 | 2.029 | 0.042 |
| Tracheophyta | Species trait | Range size | 0.48 | 0.129 | 0.226 | 0.733 | 3.714 | <0.001 |
| Tracheophyta | Species trait | Domain [terrestrial] | NA | NA | NA | NA | NA | NA |
| Tracheophyta | Species trait | IUCN [threatened] | NA | NA | NA | NA | NA | NA |
| Tracheophyta | Species trait | Domain [terrestrial] | 1.363 | 0.921 | -0.442 | 3.168 | 1.48 | 0.139 |
| Tracheophyta | Cultural trait | IUCN [threatened] | 2.218 | 0.558 | 1.124 | 3.313 | 3.975 | <0.001 |
| Tracheophyta | Cultural trait | IUCN [non-threatened] | 1.633 | 0.433 | 0.785 | 2.482 | 3.772 | <0.001 |
| Tracheophyta | Species trait | Taxonomic uniqueness (Genus) | -0.02 | 0.114 | -0.244 | 0.204 | -0.177 | 0.859 |
| Tracheophyta | Cultural trait | Common name [yes] | 1.37 | 0.326 | 0.731 | 2.008 | 4.203 | <0.001 |
| Tracheophyta | Cultural trait | Human use [yes] | 0.985 | 0.281 | 0.435 | 1.535 | 3.508 | <0.001 |
| Tracheophyta | Cultural trait | Harmful to humans [yes] | -0.169 | 0.549 | -1.244 | 0.906 | -0.308 | 0.758 |
