## Appendix 1 for "Drivers of species knowledge across the Tree of Life"

**Appendix 2**

### Drivers of species knowledge across the Tree of Life

*Mammola S., et al.*

#### ADDITIONAL TEXT

Information on trait extraction for the different groups. Note that the grouping of taxa is not always at the same level of Linnaean rank, but rather it reflects the groups that were tackled by the different experts (see author list).

**Acanthocephala.** We derived data on specific traits from the original description (or redescription) of each. Average size is the total body length. Information on size according to sex is unknown or not reported for most taxa.

**Annelida.** We derived data on specific traits from the original description (or redescription) of each. Average size is the total body length. Information on size according to sex is unknown or not reported for most taxa.

##### Arthropoda

*Arachnida.* For Araneae, we derived specific traits from the original description of each species (World Spider Catalog, 2022), and complemented information by searching relevant papers and information in Google and Google Scholar. Male and female size is the total body length (prosoma + opisthosoma), including chelicerae. Habitat is simply the habitat of the type locality unless additional information was available in ecological/faunistic papers. For other arachnid orders, we derived specific traits from the original descriptions of the species (World Arachnida Catalog; https://wac.nmbe.ch/), and complemented information by searching relevant papers and information in Google and Google Scholar. For Acari, total body length is given after excluding the gnathosoma. For oribatid mites, total body size is given as the length of notogaster + prodorsum. For classes with a tail or telson (e.g., scorpions and microwhip scorpions), body length is the sum of prosoma + opisthosoma + tail/telson.

*Insecta.* We retrieved the imaginal phase trait information by searching relevant papers and information in Google, Google Scholar, the Biodiversity Heritage Library and in entomological books. Male and female size is the total body length (head + thorax + abdomen). We reported only the mean body length when no information about the sex was available. We classified the domain as the natural environment of the type locality unless additional information was available in supplementary literature.

*Merostomata.* We derived specific traits from the original description of each species when available and from additional relevant literature, including taxonomic reviews and monographs. Information on size according to sex is unknown or not reported for most taxa. Habitat corresponds to that of the type locality unless additional information was available in the supplementary literature.

*Myriapoda.* Specific traits for each species were derived from original descriptions and/or additional relevant literature. The majority of the information about the literature was found on Millibase (https://www.millibase.org/aphia.php?p=search). Body size is the distance between the anterior margin of the head and the posterior margin of the last body segment, excluding the antennae and/or the last pair of legs. Original descriptions and/or additional relevant literature were used to extract habitat information.

*Entognatha.* We derived specific traits from the original description of each species when available and from additional relevant literature, including taxonomic reviews and monographs. Information on size according to sex is unknown or not reported for most taxa. Habitat corresponds to that of the type locality unless additional information was available in the supplementary literature.

*Pygnogonida.* We derived specific traits from the original description of each species when available and from additional relevant literature, including taxonomic reviews and monographs. Information on size according to sex is unknown or not reported for most taxa. Habitat corresponds to that of the type locality unless additional information was available in the supplementary literature.

**Basidiomycota.** We retrieved data through web-based searches using the Google search engine. When available, we assessed the original species description using the Biodiversity Heritage Library (https://www.biodiversitylibrary.org/). We retrieved habitat information from online searches and compared it with GBIF data. We ranked species as harmful if known to be poisonous or toxic to humans. We considered the pileus/fruiting body diameter as the average size.

**Bryozoa.** We derived data on specific traits from the original description of each species when available and from additional relevant literature, including taxonomic reviews, monographs, and online databases (WORMS; Horton and et al., 2021). Average size is the mean zooid length. Information on size according to sex is unknown or not reported for most taxa. Habitat corresponds to that of the type locality unless additional information was available in the supplementary literature.

**Chaetognata.** We derived data on specific traits from the original description of each species (when available) and from additional relevant literature, including taxonomic reviews, monographs, and online databases (WORMS; Horton and et al., 2021). Average size is the total body length. Information on size according to sex is unknown or not reported for most taxa. Habitat corresponds to that of the type locality unless additional information was available in the supplementary literature.

**Chordata**

*Amphibia.* We retrieved the average size and domain for most species from Oliveira et. al. (2017). The remaining length values were mostly taken from the database AmphibiaWeb, either English or French language accounts, and very few had to be retrieved from the original description or other literature. We considered Amphibian species as harmful if they were classified as invasive/pests. No venomous urodele or anuran species were present in the list. Caecilian secretion inoculation systems were only very recently described and their toxicity is still unclear, but nevertheless these animals were not considered harmful since no envenomations are known. Harm due to improper ingestion or preparation (e.g. amphibians with toxic secretions) was also not considered, since every food item has the potential to be harmful. Species solely harmful to human pets were not considered as harmful to humans.

*Aves.* For each species, we used data from various online sources, especially oiseaux.net, birdsoftheworld.org and the IUCN Red List (IUCN, 2020) to find specific traits. We derived additional information from various books (Cramp and Perrins, 1994; Sibley, 2000; Stevenson and Fanshawe, 2002), and from scientific articles located through searches in *Web of Science*. Details of the measurements of body size were rarely given, but the standard measurement is bill tip to tail tip of a laid-out specimen, which we assumed unless specified otherwise. Information on human use, domain, habitat and trophic role was based primarily on that given in the IUCN Red List (IUCN, 2020); otherwise, we used scientific publications. We selected red or blue colorations based on 'strikingness' rather than extent (e.g. a black bird with a small bright red crown would be scored as red colored) assessed from online images (images were available for all selected bird species).

*Fish (Actinopterygii, Chondrichthyes, Hyperoartia, Myxini and Sarcopterigii).* We collected data on fish species from these five classes from several online databases such as Fishbase, the IUCN red list (IUCN, 2020), and WORMS (Horton and et al., 2021). We gathered data on domain, reproductive habitat and trophic role in the specialized literature when not available on the online databases. Body size refers to the maximum body length. The field "human use" includes both species used for human consumption and popular species for aquarists. In the latter case, we checked specialized webpages to determine if a species was regularly used as an aquarium fish.

*Mammalia.* We compiled data on mammals from a variety of sources including the Handbook of Mammals of the World (body size measurements, Diet), AnimalDiversity web (body size measurements, human use), and the IUCN Red List (IUCN, 2020) archive (habitat preferences and human use). We expressed body size as head-body length in all cases except cetaceans for which size measures are expressed as total length (including the tail length).

*Reptilia.* We retrieved squamate body length (maximum SVL) retrieved from Feldman et al. (2019). Data comprise SVL for lizards and amphisbaenians, while for snakes most of the data (c. 90%) comprise total lengths (TL) and the rest SVL. Crocodilian body length values were obtained from Trutnau and Sommerlad (2006). Turtle carapace lengths were obtained from Itescu et al. (2014). It should be noted that there is a reptile-standardized mass size-value, based on clade-specific allometric equations [updated in Slavenko et al. (2019, 2016)], but that was not used to keep coherence with other taxa in this study.

**Cnidaria.** We derived data on specific traits from the original description of each species when available and from additional relevant literature, including taxonomic reviews, monographs and online databases (WORMS; Horton and et al., 2021). Average size is the total body length of a specimen. The size of a colonial species correlates to the overall size of the colony since this is what is recognised in the field. Information on size according to sex is unknown or not reported for most taxa. Habitat corresponds to that of the type locality or to the habitat stated in the review monographs unless additional information was available in the supplementary literature.

**Ctenophora.** We derived data on specific traits from the original description of each species when available and from additional relevant literature, including taxonomic reviews and monographs (WORMS; Horton and et al., 2021). Average size is the body length measured from the aboral pole to the tip of the mouth. Information on size according to sex is unknown or not reported for most taxa. Habitat corresponds to that of the type locality unless additional information was available in the supplementary literature.

**Cycliophora.** We derived data on specific traits from the original description of each species and from additional relevant literature. Average size is the total body length. Information on size according to sex is unknown or not reported. Habitat corresponds to that of the type locality unless additional information was available in the supplementary literature.

**Echinodermata.** We derived specific traits from the original description of each species when available and from additional relevant literature, including taxonomic reviews, monographs and online databases (WORMS; Horton and et al., 2021). Average size is body length for Holothuroidea, body diameter for Echinoidea and Asteroidea, arm length for Crinoidea, and disc diameter measured from the distal edge of the radial shields to the edge of the opposite interradial for Ophiuroidea. Information on size according to sex is unknown or not reported for most taxa. Habitat corresponds to that of the type locality unless additional information was available in the supplementary literature.

**Hemichordata.** We derived data on specific traits from the original description of each species when available and from additional relevant literature, including taxonomic reviews, monographs and online databases (WORMS; Horton and et al., 2021). Average size is the total body length. Information on size according to sex is unknown or not reported for most taxa. Habitat corresponds to that of the type locality unless additional information was available in the supplementary literature.

**Kinorhyncha.** We derived data on specific traits from the original description of each species when available and from additional relevant literature, including taxonomic reviews, monographs and online databases (WORMS; Horton and et al., 2021) Average size is the total body length. Information on size according to sex is unknown or not reported for most taxa. Habitat corresponds to that of the type locality unless additional information was available in the supplementary literature.

**Loricifera.** We derived data on specific traits from the original description of each species when available and from additional relevant literature, including taxonomic reviews, monographs and online databases (WORMS; Horton and et al., 2021). Average size is the body length excluding the mouth cone. Information on size according to sex is unknown or not reported for most taxa. Habitat corresponds to that of the type locality unless additional information was available in the supplementary literature.

**Mollusca.** We derived data on specific traits from the original description of each species when available and from additional relevant literature, including taxonomic reviews, monographs and online databases (MolluscaBase; MolluscaBase eds., n.d.). Average size is the total body length. Information on size according to sex is unknown or not reported for most taxa. Habitat corresponds to that of the type locality unless additional information was available in the supplementary literature.

**Nemertea.** We derived specific traits from the original description of each species when available and from additional relevant literature, including taxonomic reviews, monographs and online databases (WORMS; Horton and et al., 2021). Average size is the total body length. Information on size according to sex is unknown or not reported for most taxa. Habitat corresponds to that of the type locality unless additional information was available in the supplementary literature.

**Nematoda.** We derived data on specific traits from the original description of each species when available and from additional relevant literature, including taxonomic reviews and monographs. Average size is the total body length. Information on size according to sex is unknown or not reported for most taxa. Habitat corresponds to that of the type locality or to the habitat stated in the review monographs unless additional information was available in the supplementary literature

**Nematomorpha.** We derived specific traits from the original description of each species when available and from additional relevant literature, including taxonomic reviews, monographs and online databases (WORMS; Horton and et al., 2021). Average size is the total body length. Information on size according to sex is unknown or not reported for most taxa. Habitat corresponds to that of the type locality unless additional information was available in the supplementary literature.

**Platyhelminthes.** We derived data on specific traits from the original description of each species when available and from additional relevant literature, including taxonomic reviews, monographs and online databases (Gibson et al., 2014; Horton and et al., 2021). Average size is the total body length. Information on size according to sex is unknown or not reported for most taxa. Habitat for parasitic species corresponds either to the habitat of the animal they parasite or (when known) the domain where the larval free-living stages inhabit.

**Porifera.** We derived specific traits from the original description of each species when available and from additional relevant literature, including taxonomic reviews, monographs and online databases (WORMS; Horton and et al., 2021). Average size is the total body length of a specimen. Information on size according to sex is unknown or not reported for most taxa. Habitat corresponds to that of the type locality or to the habitat stated in the review monographs unless additional information was available in the supplementary literature.

**Priapulida.** We derived specific traits from the original description of each species when available and from additional relevant literature, including taxonomic reviews, monographs and online databases (WORMS; Horton and et al., 2021). Average size is the body length excluding the tail. Information on size according to sex is unknown or not reported for most taxa. Habitat corresponds to that of the type locality unless additional information was available in the supplementary literature.

**Rotifera.** We derived specific traits from the original descriptions and published literature, especially guides and taxonomic reviews (extracted from Jersabek and Leitner, 2013). Size is the total body length when fully extended, which is not available for the yet unknown males for most species.

**Tardigrada.** We derived specific traits from the original description of each species when available and from additional relevant literature, including taxonomic reviews, monographs and online databases (WORMS; Horton and et al., 2021). Average size is the total body length. Information on size according to sex is unknown or not reported for most taxa. Habitat corresponds to that of the type locality unless additional information was available in the supplementary literature.

**Viridiplantae (Anthocerotophyta, Bryophyta, Marchantiophyta, Tracheophyta) and Rhodophyta.** We retrieved data from web-based searches, using the species' binomial in the Google search engine. Where available, we obtained data from online databases and websites encompassing the main floras worldwide. If no information was retrieved, we accessed the original species description using the Biodiversity Heritage Library (https://www.biodiversitylibrary.org/) (if available). Given the differences in taxonomies adopted in GBIF and the different mined sources, when searches using GBIF taxonomy did not return any useful data, we used synonyms. We retrieved habitat information from accessed sources (both web-based searches and literature) when available, and compared with available information in GBIF. We checked species usefulness for humans using the 'Useful Tropical Plant' (http://tropical.theferns.info/) and the 'Pl@ntUse' (https://uses.plantnet-project.org/) databases or by web searches looking for uses as raw materials, food or other purposes such as cosmetics, medicine, sources of phytochemicals or domesticated plants. We classified species as harmful if they are known to be poisonous or toxic to humans. We calculated the average size using the maximum and minimum stem height for herbs, or overall size for trees and shrubs. When not available, we only considered the maximum size. For algae and bryophytes, we approximated size as the thallus height.
